# Running with an Exotendon Reduces Compressive Knee Contact Force

**DOI:** 10.1101/2025.11.19.689319

**Authors:** Jon Stingel, Nicos Haralabidis, Jennifer Hicks, Scott Uhlrich, Scott Delp

## Abstract

An exotendon—a spring that couples the dynamics of the legs when attached to a runner’s shoes—reduces the energetic cost of running, but the effects on joint contact forces are unknown. This study examined whether running with an exotendon alters the forces in the hip, knee and ankle. We used muscle-driven simulations of experimental data to compute compressive and shear contact forces at the hip, knee, and ankle joints for five participants running at 2.7 m/s with and without an exotendon. We found that runners using the exotendon experienced a 9.4% reduction in peak knee compressive contact force (1.0 ± 0.6 BW; *P*=0.036), and no change in the peak knee shear contact force. The primary contributor to this reduction was lower forces in the quadriceps muscles, which decreased their contribution to peak knee compressive contact force by 14.2% (-0.9 ± 0.6 BW; P=0.026). We observed no change in the peak compressive or shear contact forces in the hip or ankle joints. Though the exotendon was not originally designed to reduce joint forces, our findings highlight the ability of this simple device to make changes to gait that reduce both energetic cost and compressive knee force.

## 1. Introduction

Exercise is vital for maintaining health. It preserves muscle strength and volume (Wisdom et al., 2015), supports skeletal tissue integrity (Frost, 1994), improves cardiopulmonary function, and contributes to cartilage and connective tissue health (Bader et al., 2011; Sun, 2010). Running is one of the most popular forms of exercise, and emerging evidence suggests that moderate-intensity exercise, such as running, supports tissue health (Bramble and Lieberman, 2004; Halilaj et al., 2018; Lai et al., 2014; Wakimoto et al., 2024). For example, athletes who regularly load their joints demonstrate increased knee cartilage thickness and proteoglycan content (Gratzke et al. 2007; Helminen 2009; Tiderius et al. 2004).

Excessive loading, rapid increases in training intensity, or abrupt changes in loading patterns, however, can accelerate tissue degradation and lead to injury (Andriacchi et al., 2009; Frandsen et al., 2025; Lin et al., 2013). Reducing knee joint forces is typically the focus of interventions aimed at mitigating knee pain and injury risk during gait (Alexander et al., 2022; Uhlrich et al., 2022; Wan et al., 2024). Compressive joint force (Fig. 3) has often been studied since compressive forces in the knee can reach between 7-15 BW in activities such as running (Glitsch and Baumann, 1997; Miller et al., 2014; Rooney and Derrick, 2013; Saxby et al., 2016), but shear force is also important in understanding joint health (Carter et al., 2004; Wilson et al., 2003).

An exotendon is a passive assistive device that reduces runners’ energy expenditure (Stingel et al., 2023). Since the exotendon lowers knee extension and hip flexion moments and the rate of energy expenditure (Simpson et al., 2019), it is possible that the joint contact forces are also reduced. Muscle forces account for more than half of the knee joint contact force during gait (Sasaki and Neptune, 2010; Uhlrich et al., 2022). Runners who reduced their rate of energy expenditure from using the exotendon also saved energy in their quadriceps muscles, by decreasing muscle activation and muscle force (Stingel et al., 2023). Energetic savings and reductions in muscle forces were also observed in hip-spanning muscles, which could decrease forces at the hip joint. Conversely, no changes were detected in ankle musculature and therefore the ankle joint may not exhibit any differences in force.

This work examines how joint contact forces change when running with an exotendon compared to natural running. Based on the previously observed decrease in knee extension moments and quadriceps activations, we hypothesized that running with an exotendon would reduce the peak compressive and shear knee contact forces, and that the contribution of quadriceps force to compressive forces at the knee joint would decrease. In addition, we explored whether there were changes in the peak hip and ankle compressive and shear contact forces.

## 2. Materials and methods

### 2.1 Experimental Data

We used previously collected experimental data of five healthy recreational runners (age: 26 ± 1 year; height: 170.9 ± 9.2 cm; mass: 63.5 ± 8.1 kg) who ran on a treadmill at 2.7 m/s with and without the exotendon (Simpson et al., 2019; Stingel et al., 2023). The study was approved by the Stanford University Institutional Review Board, and all participants provided written informed consent prior to participation. Prior to data collection, participants completed two training sessions. In each session, they ran four 10-minute trials on a treadmill alternating between exotendon and natural to enable them to adapt to running on the treadmill and with the exotendon. On the data collection day, participants ran on the same instrumented treadmill, which measured three-dimensional (3D) ground reaction forces (Bertec Corporation, Columbus, OH, USA). They completed a 7-minute run with and without the exotendon, with a 5-minute break between each condition in randomized order. Prior to the runs, participants were fitted with 40 retro-reflective markers for 3D, full-body motion capture (Motion Analysis Corporation, Santa Rosa, CA, USA). Data from the final 2 minutes of each trial was analyzed to ensure that each runner had achieved a steady gait pattern.

### 2.2 Musculoskeletal model

We used a musculoskeletal model in OpenSim 4.5 (Seth et al., 2018) with 37 degrees of freedom, 80 muscle-tendon units that spanned the lower limbs, and idealized torque generators that actuated the trunk and upper limbs (Arnold et al., 2010; Rajagopal et al., 2016; Uhlrich et al., 2022). The generic model was scaled to each participant’s anthropometry using a standing static trial; the maximum isometric force for each muscle was adjusted by scaling the total muscle volume for each participant based on their mass and height (Handsfield et al., 2014). The exotendon was modeled as a linear spring connecting the calcanei at measured locations during the experiment (Fig. 1). Using the adjusted models, we performed an inverse kinematics analysis to compute the joint angles and positions for 4 gait cycles in each condition. We filtered the kinematics and measured ground reaction forces at 15 Hz with a fourth order lowpass Butterworth filter and used them as input for an inverse dynamics analysis where we computed the joint moments during each of the gait cycles.

**Fig. 1.**
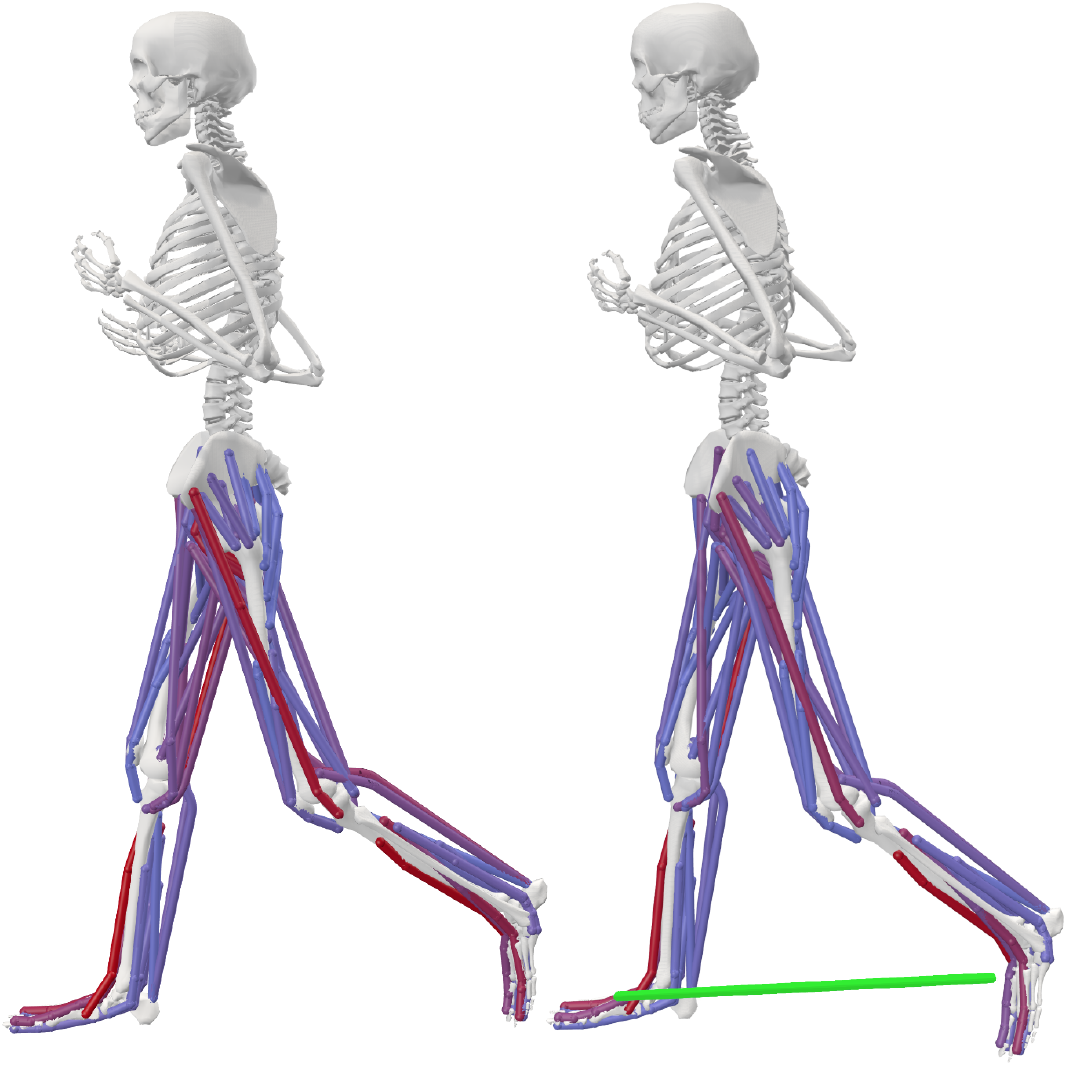
Natural (left) and exotendon (right) versions of OpenSim models (Arnold et al., 2010; Rajagopal et al., 2016; Uhlrich et al., 2022) used to simulate experimental runners. The exotendon (green) was modeled as a spring and connected the runner’s feet.

### 2.3 Optimal control simulations

We generated muscle-driven kinematic tracking simulations of each participant and each condition using OpenSim Moco (Dembia et al., 2020). To create these simulations, we provided as input the inverse kinematics joint angles and positions, inverse dynamics joint moments, scaled musculoskeletal model, and 3D ground reaction forces and moments. The simulation solves for a set of joint angles, positions, and moments that satisfy the dynamics of the prescribed input ground reaction forces and moments, as well as the muscle states (muscle activations and fiber lengths) required to generate the experimentally measured motion. Reserve actuators (Delp et al., 2007; Hicks et al., 2015) with an optimal force of 1 Nm were added to the model to ensure we could generate the input motions, but their usage was penalized in the cost function to encourage the optimizer to use the simulated muscles to generate the experimentally observed movement dynamics. The highest weighted term in the optimization cost function minimized deviation from the input joint angles and positions. We also minimized the sum of muscle excitations and torque actuator controls, and the deviation from input joint moments. This resulted in a dynamically consistent simulation with prescribed ground reaction forces that tracked inverse kinematics and inverse dynamics results.

We previously used a similar musculoskeletal model and optimal control framework to examine muscle contributions to metabolic cost in natural and exotendon running (Stingel et al., 2023), and found that the framework captured experimentally measured reductions in whole-body metabolic cost and produced between-condition changes in muscle activity that aligned with electromyography. These results provide confidence in the framework’s ability to predict changes in joint contact forces between conditions, which are largely driven by muscle forces.

For each runner, we simulated a total of eight gait cycles, split between natural and exotendon running, and used OpenSim’s Joint Reaction Analysis tool to compute the resultant forces at each joint due to the muscles, external forces, and internal forces of the model (Steele et al., 2012). This allowed us to examine resultant joint contact forces, contact forces acting in joint axial directions, and contact forces due to specific muscle contributions. We performed the analysis for the hip, knee, and ankle joints. We then segmented the resultant contact forces into compressive and shear contact components at each joint, where the compressive component is the force directed along the axial direction of the distal bone in the joint and the shear is the transverse component of the force in the joint. All data and code from these simulations are available online at https://github.com/stingjp/exotendonJointContactPaper.git.

### 2.4 Statistical analyses

To reduce the chance of Type I error (i.e., to control the familywise error rate), we used a fixed-sequence hierarchical testing procedure (Dmitrienko and Tamhane, 2007). We defined three tiers of statistical tests, with each subsequent tier evaluated only if at least one comparison in the preceding tier reached significance. For each tier of tests, we used a significance level of α = 0.05. Within each tier, we used a Bonferroni correction for multiple comparisons, and we reported corrected p-values.

For the first tier, we hypothesized that compressive and shear knee contact forces would be reduced with an exotendon. Peak compressive and shear knee contact forces were compared between conditions using two Bonferroni-corrected two-tailed Student’s t-tests. For the second tier, we hypothesized that the quadriceps contribution to the peak compressive knee contact force would be reduced with an exotendon. This was tested using a two-tailed Student’s t-test. For the third tier, we explored differences in the compressive and shear forces in the hip and ankle joints. Peak compressive and shear forces in both the hip and ankle joints were compared between conditions using four Bonferroni-corrected two-tailed Student’s t-tests.

## 3. Results

Peak compressive knee contact force decreased by 9.4 ± 4.7% (1.0 ± 0.6 BW; *P*=0.036) (Fig. 2a) when running with an exotendon. There was no change detected in peak shear knee contact force between exotendon and natural running (4.2 ± 14.7%; 0.04 ± 0.2 BW; *P*=1.0) (Fig. 2b).

**Fig. 2.**
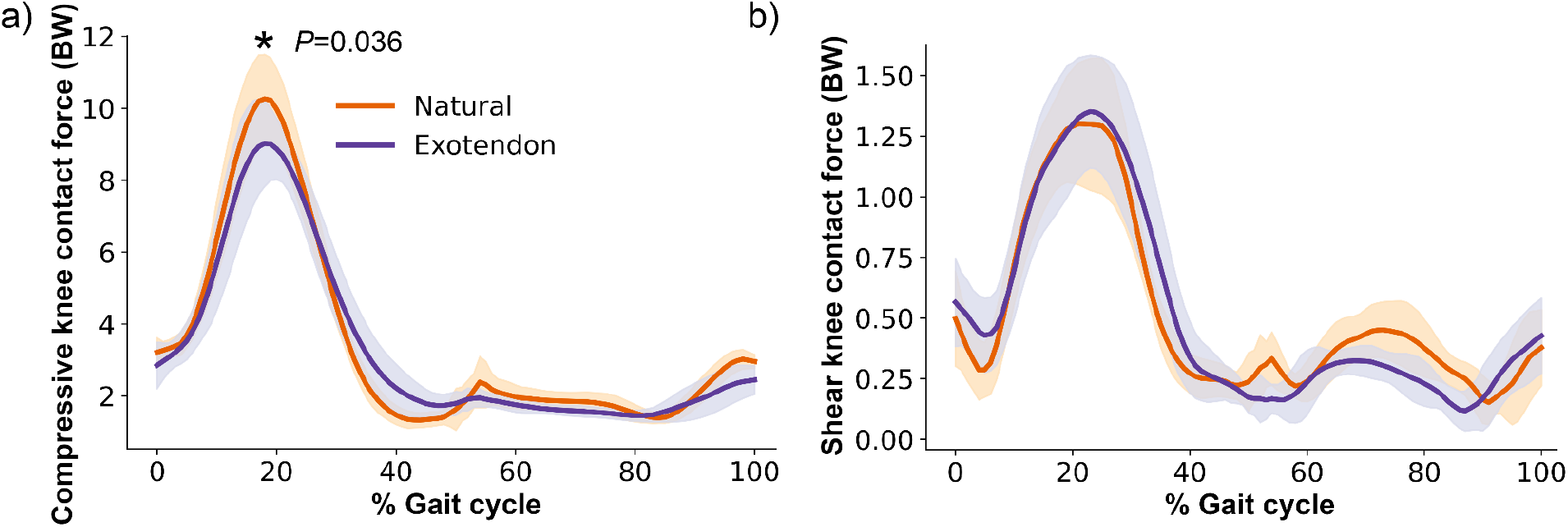
The average a) compressive and b) shear knee contact forces are shown throughout the gait cycle for natural (orange) and exotendon (purple) running. The solid line indicates the average across 5 participants, 4 gait cycles, and both legs; the shaded regions indicate the standard deviation. ^*^ indicates a statistically significant difference in peak values using a paired student’s t-test with a Bonferroni two-comparison correction.

**Fig. 3.**
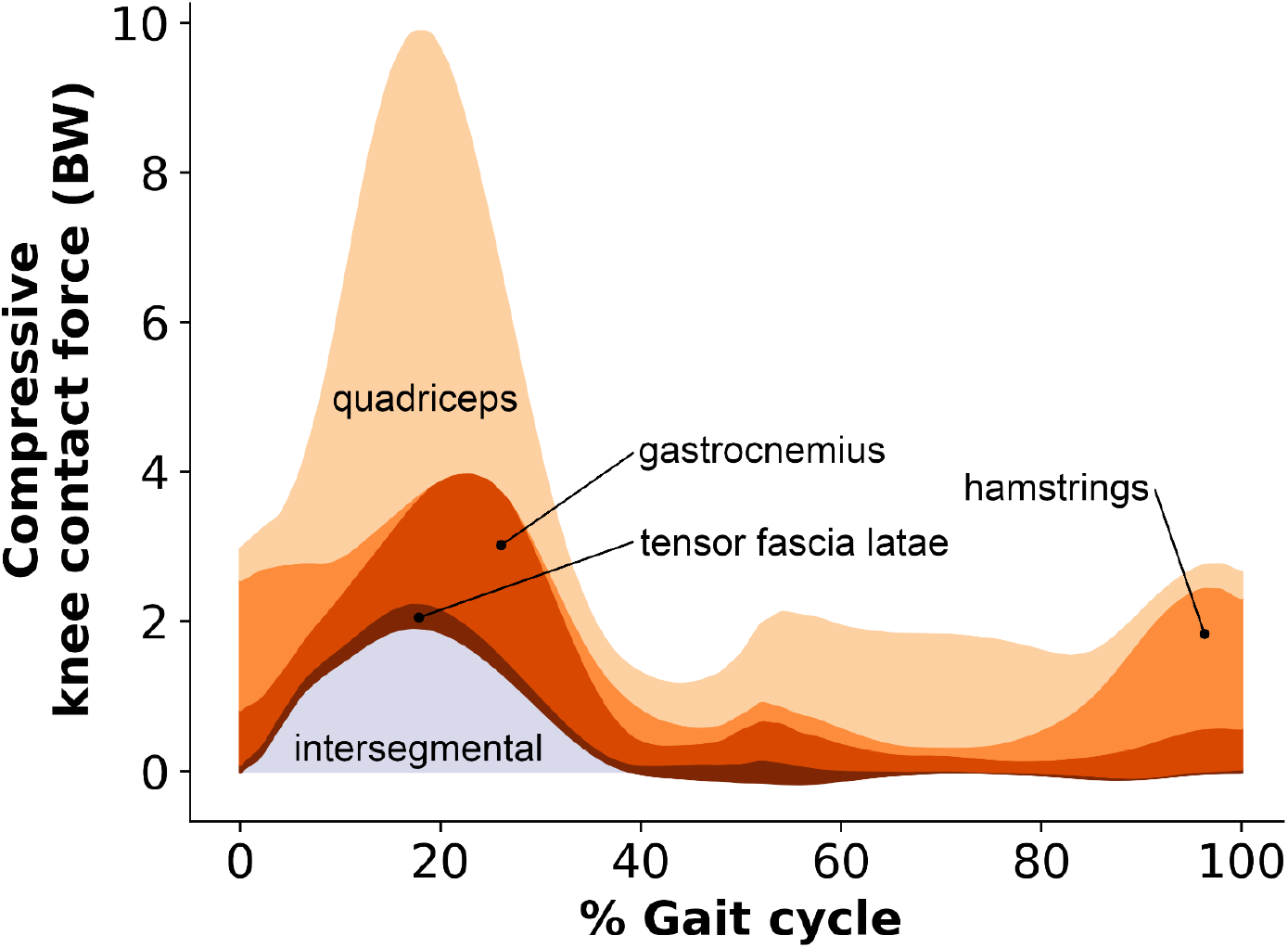
Contributions to compressive knee contact force during natural running at 2.7 m/s. The contributions of the intersegmental force (purple) and muscle forces (orange) to compressive knee contact force are shown, such that each force contributes cumulatively to the total compressive knee contact force. Contact forces are expressed in terms of bodyweight (BW) and averaged for the 5 subjects included in the current study.

The quadriceps were the largest contributor to the stance-phase peak in compressive knee contact force during natural running, accounting for 60% of the peak compressive force (Fig. 3). The quadriceps also showed the largest change in contact force contributions. The contribution of the quadriceps to peak compressive knee contact force was reduced when running with an exotendon by 14.2 ± 8.7% (0.9 ± 0.6 BW; *P*=0.026) (Fig. 4) compared to natural running.

**Fig. 4.**
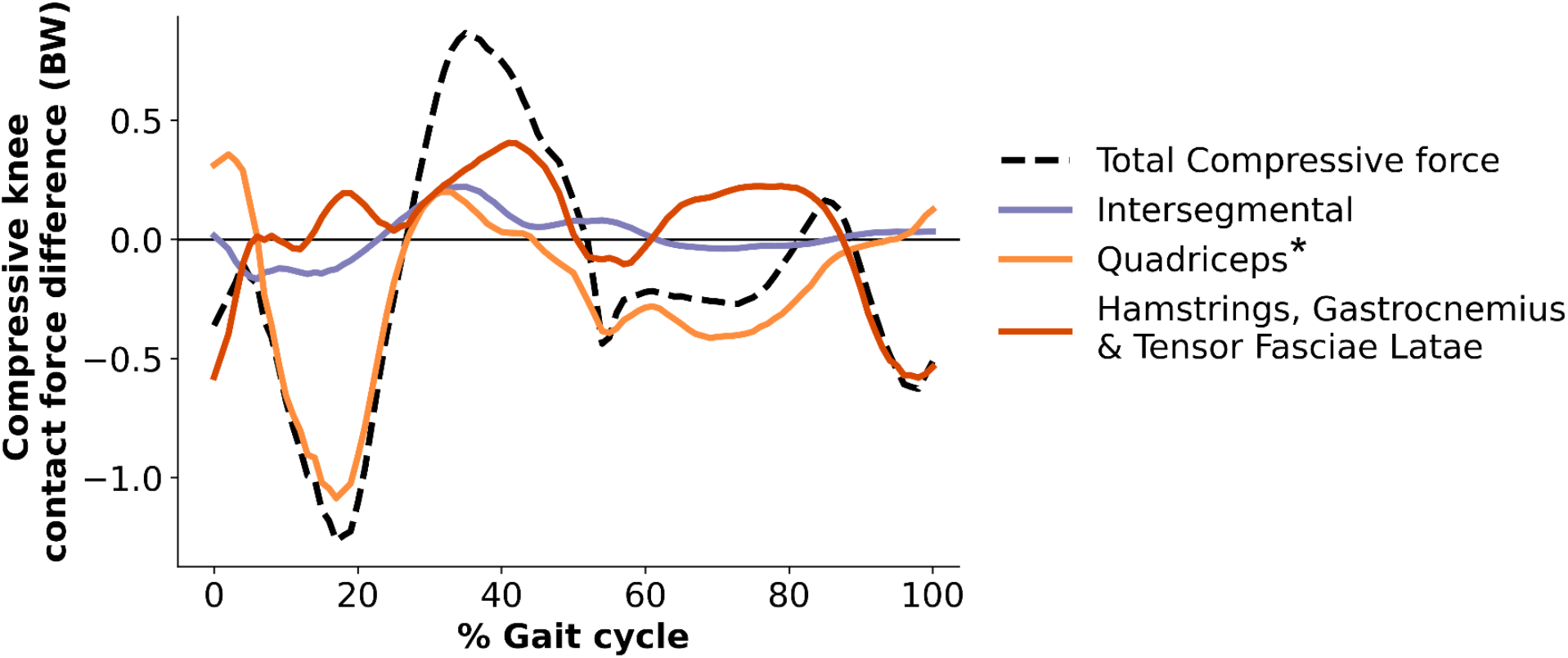
Changes in compressive knee contact force from natural to exotendon running. A negative value indicates a reduction in force for exotendon running. The dashed black line is the change in total compressive knee contact force throughout the gait cycle. The changes in contributions to compressive knee contact force from the quadriceps (light orange), all other knee spanning muscles (dark orange), and the intersegmental force (purple line) are also plotted. Each line represents the average across five participants, 4 gait cycles, and both legs. ^*^ indicates a statistically significant difference in peak values using a paired student’s t-test with a significance level of α=0.05.

We detected no changes in the peak compressive hip contact force (-1.5 ± 10%; -0.12 ± 0.68 BW; *P*=1.0), peak shear hip contact force (-5.6 ± 14.9%; -0.1 ± 0.4 BW; *P*=1.0), peak compressive ankle contact force (-5.3 ± 8.9%; -0.6 ± 0.9 BW; *P*=0.84), or peak shear ankle contact force (-10.0 ± 13.9%; -0.42 ± 0.6 BW; *P*=0.80) during exotendon running (Fig. 5).

**Fig. 5.**
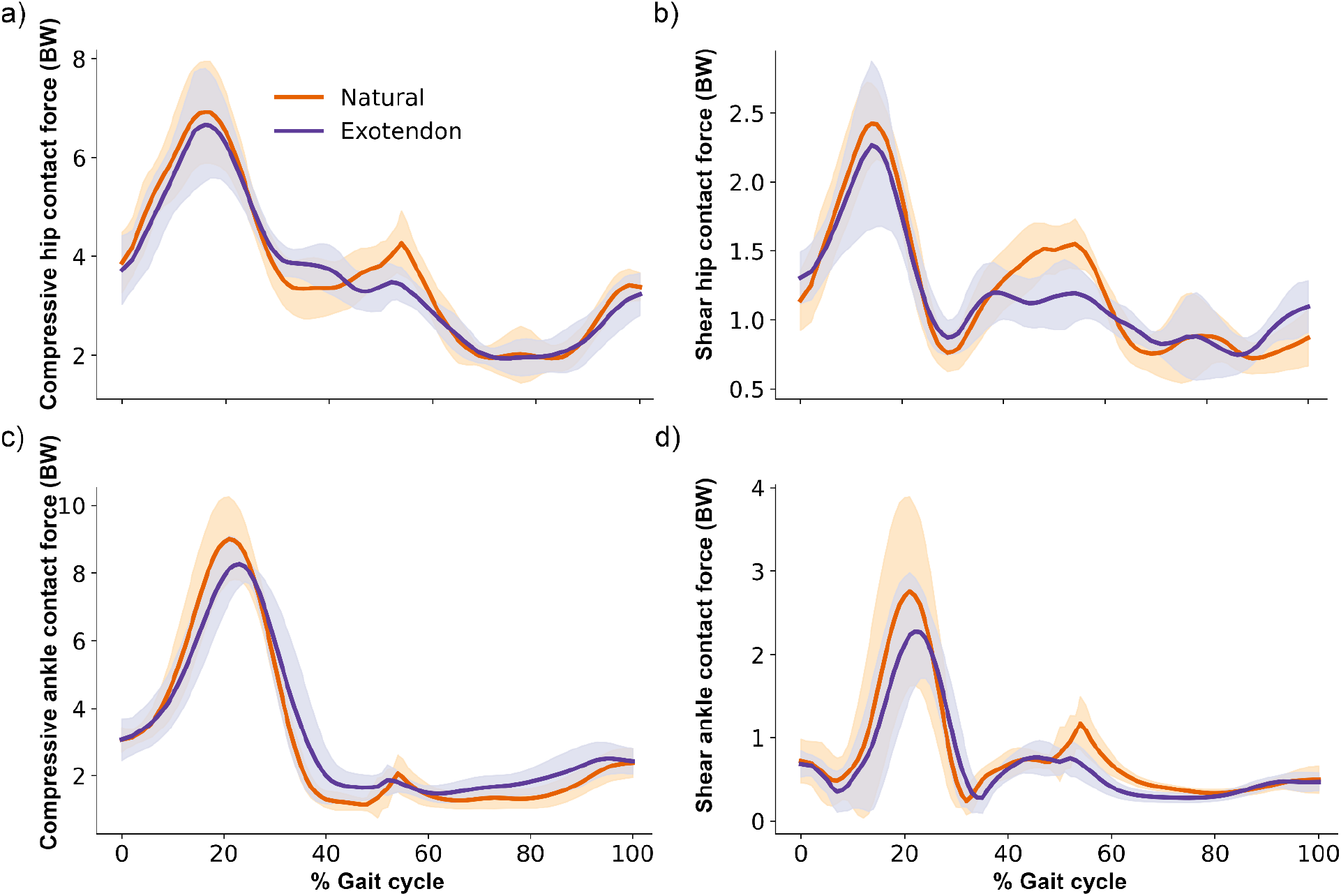
Average compressive (a, c) and shear (b, d) contact forces in the hip and ankle joints throughout the gait cycle for natural (orange) and exotendon (purple) runs. The solid line indicates the average across 5 participants, 4 gait cycles, and both legs; the shaded regions indicate the standard deviation.

## 4. Discussion

The purpose of this study was to determine whether running with an exotendon—a simple, passive assistive device—alters lower-limb joint contact forces during running. Musculoskeletal simulations revealed that peak compressive knee contact forces were significantly reduced when participants ran with the exotendon, whereas no changes were detected in other contact forces. During running at 2.7 m/s, the simple device, which has been shown to transmit approximately 30 N of peak inter-leg force (Simpson et al., 2019), facilitated a reduction in compressive knee contact force of over 600 N or about 1.0 BW on average. The reductions in knee joint force with exotendon use can be explained by changes in stride mechanics and quadriceps forces. Running with an exotendon tends to make it metabolically optimal for runners to increase stride frequency, reduce stride length, reduce knee flexion, and reduce knee extension moments (Simpson et al., 2019). Simulations confirmed these changes and showed a reduction in quadriceps activation and force in combination (Stingel et al., 2023). The current simulations also allowed us to calculate knee contact force and partition it into contributions from individual muscle groups, revealing that the reduction in peak compressive knee contact force was primarily attributable to decreased quadriceps contributions. These adaptations could be desirable for people with knee pain, as increased forces, particularly from the quadriceps, in activities of daily living can lead to worsened pain (Wan et al., 2025).

Peak knee joint contact forces were approximately 10 BW in our simulations of running at 2.7m/s. This is higher than joint forces measured with instrumented implants, which range from 3.3-7.1 BW when jogging at 1.7-1.8 m/s (Bergmann et al., 2014; Taylor and Walker, 2001). These smaller joint forces are expected at lower running speeds. Simulations of running at 2.0-3.2 m/s report knee forces of 8.0-11.3 BW (Kloock et al., 2025; McDonnell et al., 2019; Miller et al., 2014), while those at 4.0-5.0 m/s report knee forces of 7.8-15.0 BW (Edwards et al., 2008; Glitsch and Baumann, 1997; Rooney and Derrick, 2013; Saxby et al., 2016; Thomas et al., 2020). Accordingly, our simulated forces at 2.7 m/s fall within the range expected for intermediate speeds between the very slow and fast running conditions reported previously.

Our study examined joint forces in only five runners. Even with this small number of participants there was a clear and consistent reduction in quadriceps force and compressive knee loads. We did not detect changes in the shear force at the knee, or the joint forces at the hip or ankle, but we cannot conclude that the forces in these other joints do not change given the limited statistical power of our study.

The exotendon is a simple passive device that assists runners in reducing their knee contact force and lowering their rate of energy expenditure. To date, all experiments have involved healthy recreational athletes running at a single speed; thus, the possibility of reducing knee forces in other populations or at other speeds is speculative at this time and warrants further investigation. Future work should also examine whether reductions in metabolic cost and joint forces extend to individuals who may benefit from reduced knee loading, such as older adults or those who experience running-related pain.

## 5. Conclusion

We used musculoskeletal modeling and optimal control to compute joint contact forces at the hip, knee, and ankle during running with and without a passive assistive device. The simulations revealed that the device reduced compressive knee joint forces compared to natural running at the same speed. These reductions were driven by decreased quadriceps contributions to knee contact forces. Our findings demonstrate how a simple assistive device can alter the distribution of joint forces during dynamic tasks such as running.

## Acknowledgements

The authors thank Julie Muccini and Nick Bianco for their assistance in this work.

## Data Availability Statement

The data that support the findings of this study are openly available at https://github.com/stingjp/exotendonJointContactPaper.git

## Declaration of generative AI and AI-assisted technologies in manuscript preparation

During the preparation of this work the authors used OpenAI’s ChatGPT as an editing tool, using it to review already written sections of the manuscript. The authors manually reviewed the tool’s suggestions and edited the content as needed. The authors take full responsibility for the content of the published article.

## References

Alexander, J.L.N., Culvenor, A.G., Johnston, R.R.T., Ezzat, A.M., Barton, C.J., 2022. Strategies to prevent and manage running-related knee injuries: a systematic review of randomised controlled trials. Br J Sports Med 56, 1307–1319. 10.1136/bjsports-2022-105553

Andriacchi, T.P., Koo, S., Scanlan, S.F., 2009. Gait mechanics influence healthy cartilage morphology and osteoarthritis of the knee. J Bone Joint Surg Am 91 Suppl 1, 95–101. 10.2106/JBJS.H.01408

Arnold, E.M., Ward, S.R., Lieber, R.L., Delp, S.L., 2010. A Model of the Lower Limb for Analysis of Human Movement. Annals of Biomedical Engineering 38, 269–279. 10.1007/s10439-009-9852-5

Bader, D.L., Salter, D.M., Chowdhury, T.T., 2011. Biomechanical Influence of Cartilage Homeostasis in Health and Disease. Arthritis 2011, 979032. 10.1155/2011/979032

Bergmann, G., Bender, A., Graichen, F., Dymke, J., Rohlmann, A., Trepczynski, A., Heller, M.O., Kutzner, I., 2014. Standardized Loads Acting in Knee Implants. PLOS ONE 9, e86035. 10.1371/journal.pone.0086035

Bramble, D.M., Lieberman, D.E., 2004. Endurance running and the evolution of Homo. Nature 2004 432:7015 432, 345–352. 10.1038/nature03052

Carter, D.R., Beaupré, G.S., Wong, M., Smith, R.L., Andriacchi, T.P., Schurman, D.J., 2004. The Mechanobiology of Articular Cartilage Development and Degeneration. Clinical Orthopaedics and Related Research® 427, S69. 10.1097/01.blo.0000144970.05107.7e

Delp, S.L., Anderson, F.C., Arnold, A.S., Loan, P., Habib, A., John, C.T., Guendelman, E., Thelen, D.G., 2007. OpenSim: Open-Source Software to Create and Analyze Dynamic Simulations of Movement. IEEE Transactions on Biomedical Engineering 54, 1940–1950. 10.1109/TBME.2007.901024

Dembia, C.L., Bianco, N.A., Falisse, A., Hicks, J.L., Delp, S.L., 2020. OpenSim Moco: Musculoskeletal optimal control. PLOS Computational Biology 16, e1008493. 10.1371/journal.pcbi.1008493

Dmitrienko, A., Tamhane, A.C., 2007. Gatekeeping procedures with clinical trial applications. Pharmaceutical Statistics 6, 171–180. 10.1002/pst.291

Edwards, W.B., Gillette, J.C., Thomas, J.M., Derrick, T.R., 2008. Internal femoral forces and moments during running: Implications for stress fracture development. Clinical Biomechanics 23, 1269–1278. 10.1016/j.clinbiomech.2008.06.011

Frandsen, J.S.B., Hulme, A., Parner, E.T., Møller, M., Lindman, I., Abrahamson, J., Simonsen, N.S., Jacobsen, J.S., Ramskov, D., Skejø, S., Malisoux, L., Bertelsen, M.L., Nielsen, R.O., 2025. How much running is too much? Identifying high-risk running sessions in a 5200-person cohort study. Br J Sports Med. 10.1136/bjsports-2024-109380

Frost, H.M., 1994. Wolff’s Law and bone’s structural adaptations to mechanical usage: an overview for clinicians. Angle Orthod 64, 175–188. 10.1043/0003-3219(1994)064%253C0175:WLABSA%253E2.0.CO;2

Glitsch, U., Baumann, W., 1997. The three-dimensional determination of internal loads in the lower extremity. Journal of Biomechanics 30, 1123–1131. 10.1016/S0021-9290(97)00089-4

Halilaj, E., Hastie, T.J., Gold, G.E., Delp, S.L., 2018. Physical activity is associated with changes in knee cartilage microstructure. Osteoarthritis and Cartilage 26, 770–774. 10.1016/j.joca.2018.03.009

Handsfield, G.G., Meyer, C.H., Hart, J.M., Abel, M.F., Blemker, S.S., 2014. Relationships of 35 lower limb muscles to height and body mass quantified using MRI. Journal of Biomechanics 47, 631–638. 10.1016/j.jbiomech.2013.12.002

Heiderscheit, B.C., Chumanov, E.S., Michalski, M.P., Wille, C.M., Ryan, M.B., 2011. Effects of Step Rate Manipulation on Joint Mechanics during Running. Med Sci Sports Exerc 43, 296–302. 10.1249/MSS.0b013e3181ebedf4

Hicks, J.L., Uchida, T.K., Seth, A., Rajagopal, A., Delp, S.L., 2015. Is My Model Good Enough? Best Practices for Verification and Validation of Musculoskeletal Models and Simulations of Movement. J Biomech Eng 137. 10.1115/1.4029304

Kloock, L., Arensmann, A., de Graaf, M.L., Gerlach, M., Boström, K.J., Wagner, H., 2025. Joint contact forces during barefoot, minimal and conventional shod running are highly individual. Sci Rep 15, 25022. 10.1038/s41598-025-09174-w

Lai, A., Schache, A.G., Lin, Y.-C., Pandy, M.G., 2014. Tendon elastic strain energy in the human ankle plantar-flexors and its role with increased running speed. J Exp Biol 217, 3159–3168. 10.1242/jeb.100826

Lin, W., Alizai, H., Joseph, G.B., Srikhum, W., Nevitt, M.C., Lynch, J.A., McCulloch, C.E., Link, T.M., 2013. Physical activity in relation to knee cartilage T2 progression measured with 3 T MRI over a period of 4 years: data from the Osteoarthritis Initiative. Osteoarthritis and Cartilage 21, 1558–1566. 10.1016/j.joca.2013.06.022

McDonnell, J., Zwetsloot, K.A., Houmard, J., DeVita, P., 2019. Skipping has lower knee joint contact forces and higher metabolic cost compared to running. Gait & Posture 70, 414–419. 10.1016/j.gaitpost.2019.03.028

Miller, R.H., Edwards, W.B., Brandon, S.C.E., Morton, A.M., Deluzio, K.J., 2014. Why Don’t Most Runners Get Knee Osteoarthritis? A Case for Per-Unit-Distance Loads. Medicine & Science in Sports & Exercise 46, 572. 10.1249/MSS.0000000000000135

Rajagopal, A., Dembia, C., DeMers, M., Delp, D., Hicks, J., Delp, S., 2016. Full body musculoskeletal model for muscle-driven simulation of human gait. IEEE Transactions on Biomedical Engineering 63, 2068–2079. 10.1109/TBME.2016.2586891

Rooney, B.D., Derrick, T.R., 2013. Joint contact loading in forefoot and rearfoot strike patterns during running. Journal of Biomechanics 46, 2201–2206. 10.1016/J.JBIOMECH.2013.06.022

Sasaki, K., Neptune, R.R., 2010. Individual Muscle Contributions to the Axial Knee Joint Contact Force during Normal Walking. J Biomech 43, 2780–2784. 10.1016/j.jbiomech.2010.06.011

Saxby, D.J., Modenese, L., Bryant, A.L., Gerus, P., Killen, B., Fortin, K., Wrigley, T.V., Bennell, K.L., Cicuttini, F.M., Lloyd, D.G., 2016. Tibiofemoral contact forces during walking, running and sidestepping. Gait & Posture 49, 78–85. 10.1016/j.gaitpost.2016.06.014

Seth, A., Hicks, J.L., Uchida, T.K., Habib, A., Dembia, C.L., Dunne, J.J., Ong, C.F., DeMers, M.S., Rajagopal, A., Millard, M., Hamner, S.R., Arnold, E.M., Yong, J.R., Lakshmikanth, S.K., Sherman, M.A., Ku, J.P., Delp, S.L., 2018. OpenSim: Simulating musculoskeletal dynamics and neuromuscular control to study human and animal movement. PLOS Computational Biology 14, e1006223. 10.1371/journal.pcbi.1006223

Simpson, C.S., Welker, C.G., Uhlrich, S.D., Sketch, S.M., Jackson, R.W., Delp, S.L., Collins, S.H., Selinger, J.C., Hawkes, E.W., 2019. Connecting the legs with a spring improves human running economy. The Journal of Experimental Biology jeb.202895. 10.1242/jeb.202895

Snyder, K.L., Farley, C.T., 2011. Energetically optimal stride frequency in running: the effects of incline and decline. J Exp Biol 214, 2089–2095. 10.1242/jeb.053157

Steele, K.M., DeMers, M.S., Schwartz, M.H., Delp, S.L., 2012. Compressive tibiofemoral force during crouch gait. Gait & Posture 35, 556–560. 10.1016/j.gaitpost.2011.11.023

Stingel, J.P., Hicks, J.L., Uhlrich, S.D., Delp, S.L., 2023. Simulating Muscle-Level Energetic Cost Savings When Humans Run With a Passive Assistive Device. IEEE Robotics and Automation Letters 8, 6267–6274. 10.1109/LRA.2023.3303094

Sun, H.B., 2010. Mechanical loading, cartilage degradation, and arthritis. Annals of the New York Academy of Sciences 1211, 37–50. 10.1111/j.1749-6632.2010.05808.x

Taylor, S.J.G., Walker, P.S., 2001. Forces and moments telemetered from two distal femoral replacements during various activities. Journal of Biomechanics 34, 839–848. 10.1016/S0021-9290(01)00042-2

Thomas, J.M., Edwards, W.B., Derrick, T.R., 2020. Joint Contact Forces with Changes in Running Stride Length and Midsole Stiffness. J. of SCI. IN SPORT AND EXERCISE 2, 69–76. 10.1007/s42978-019-00027-3

Uhlrich, S.D., Jackson, R.W., Seth, A., Kolesar, J.A., Delp, S.L., 2022. Muscle coordination retraining inspired by musculoskeletal simulations reduces knee contact force. Sci Rep 12, 9842. 10.1038/s41598-022-13386-9

Wakimoto, Y., Miura, Y., Inoue, S., Nomura, M., Moriyama, H., 2024. Effects of different combinations of mechanical loading intensity, duration, and frequency on the articular cartilage in mice. Mol Biol Rep 51, 1–14. 10.1007/s11033-024-09762-5

Wan, Y., McGuigan, P., Bilzon, J., Wade, L., 2025. Knee loading and joint pain during daily activities in people with knee osteoarthritis: A systematic review and meta-analysis. Clinical Biomechanics 122, 106433. 10.1016/j.clinbiomech.2025.106433

Wan, Y., McGuigan, P., Bilzon, J., Wade, L., 2024. The effect of foot orientation modifications on knee joint biomechanics during daily activities in people with and without knee osteoarthritis. Clinical Biomechanics 117, 106287. 10.1016/j.clinbiomech.2024.106287

Wilson, W., van Rietbergen, B., van Donkelaar, C.C., Huiskes, R., 2003. Pathways of load-induced cartilage damage causing cartilage degeneration in the knee after meniscectomy. Journal of Biomechanics 36, 845–851. 10.1016/S0021-9290(03)00004-6

Wisdom, K.M., Delp, S.L., Kuhl, E., 2015. Use it or lose it: multiscale skeletal muscle adaptation to mechanical stimuli. Biomech Model Mechanobiol 14, 195–215. 10.1007/s10237-014-0607-3

